# Immunogenicity and protective efficacy of a *Brucella abortus* L7/L12 DNA vaccine delivered via chitosan modified PLGA nanoparticles in mice

**DOI:** 10.64898/2026.02.25.707861

**Authors:** Uddhav Chaudhari, Satyabrata Dandapat, Sivasankar Panickan, Vimal Kumar, Pradeep Mahadev Sawant

**Affiliations:** Immunology Section, ICA R-Indian Veterinary Research Institute, Izatnagar-243122, India; ICMR-National Institute for Research in Reproductive and Child Health, Mumbai-400012, India; ICMR-National Institute of Virology, Pune-411001, India; ICMR-National JALMA Institute for Leprosy and other Mycobacterial Diseases, Agra-282001, India

**Keywords:** *Brucella abortus*, L7/L12 DNA, Cationic PLGA nanoparticles, Chitosan, Mice

## Abstract

The present study evaluated the immunogenicity and protective efficacy of the chitosan (CS)-modified poly-lactide-co-glycolic acid (PLGA) nanoparticles (NPs) delivering *Brucella abortus* L7/L12 DNA in a mouse model. The NPs were prepared by solvent displacement method and characterized for size, charge, morphology, cellular uptake and cytotoxicity. The cationic CS-PLGA NPs were spherical with a mean size of ∼165 nm with a positive zeta potential (+20 mV). DNA loading efficiency of 1.2% and DNA adsorption shifted zeta potential to −45 mV. *In vitro* studies in RAW 264.7 cell line demonstrated efficient uptake of the DNA loaded cationic NPs and expression of L7/L12 protein. Intramuscular immunization of the L7/L12 DNA vaccine loaded CS-PLGA NPs elicited both humoral and cell-mediated immunity with upregulation of Th1 and Th2 cytokines along with induction of IgG antibodies in mice. IFN-γ, IL-2, and IL-4 levels were significantly (*P* < 0.001) higher than control group. The protective efficacy of the DNA loaded NPs against virulent *B. abortus* 544 infection (10^5^ CFU) was significantly higher than that of the naked DNA (*P*<0.001). These findings suggest that the CS-PLGA NPs were efficiently delivered L7/L12 DNA and exhibited adjuvant potential, conferring protection against experimental murine brucellosis.

## 1. Introduction

Brucellosis is a global public health threat and an economically important disease caused by *Brucella abortus*, an intracellular facultative pathogen that affects domestic animals and humans. In cattle, brucellosis primarily affects the reproductive system causing late-term abortion, infertility, delivery of weak calves and significant economic loss to the dairy industry. Undulant fever, arthritis and endocarditis are the major signs of the disease in humans. In India, the reported prevalence in humans ranges from 0.8-26.6% and it is mostly due to *B. abortus*. There is no licensed vaccine available for human brucellosis. However, the burden of brucellosis in cattle is mainly controlled by vaccination with live attenuated vaccines, RB51 or strain 19 and in small ruminants with strain Rev 1 (*Brucella melitensis*). However, these vaccines have certain drawbacks, as they cause abortion in pregnant animals and also the vaccine virus is pathogenic and prone to the risk of reversion to virulence, which might spread to humans through contacts. These vaccines cannot be used in humans as these strains are too virulent and have a low therapeutic index. Further, they interfere with serological diagnosis and may also be secreted in milk. Being a live vaccine, it is not recommended for use in brucellosis-free countries. Therefore, safe and efficient vaccines are required to control the disease in animals and humans. Usually, the *Brucella* organisms multiply intracellularly, especially in macrophages or reticulo-endothelial system. Protective immunity against *Brucella* infection seems to be mediated by both humoral and cellular immunity; though cell mediated immunity (CMI) is expected to play a critical role in protection, in which IFN-γ and CD8+ lymphocytes are the major effectors (Oliveira and Splitter, 1995; Skendros and Boura, 2013). IFN-γ activates macrophages which exhibit anti-*Brucella* activity (Jones and Winter, 1992). To develop better vaccines, many strategies have been attempted, but they were not much efficacious than the standard vaccines S-19. However, DNA vaccines have been used successfully for many diseases (Ulmer et al., 1993; Boyer et al., 1997) as they can induce better CMI, which might also be effective against *Brucella* infection.

The DNA vaccine is an attractive alternative to the existing live attenuated vaccines, as they provoke both humoral and cellular immune responses with many advantages like convenient to design, good safety profile and low cost of production. The ribosomal protein L7/L12 is a major antigenic component with a protective role, and the gene is conserved among *Brucella* species, which is immunodominant that induces both Th1 and Th2 mediated immune responses (Bachrach et al., 1994; Golshani et al., 2015). Therefore, immunization with such DNA vaccine along with efficient delivery system might be beneficial for induction of better immune response against *Brucella*. In this context, the use of poly-lactide-co-glycolic acid (PLGA) and chitosan nanoparticles (NPs) or microparticles for delivery of vaccine candidates has advantages because of their property of slow and pulsatile release of antigen and these are potent inducers of humoral and CMI (Silva et al., 2016; Dmour and Islam, 2022). Better immune response with caprolactone microparticles coated with *Brucella* outer membrane complex has been reported against *Brucella ovis* infection in rams (Muñoz et al., 2022). The PLGA NPs are well recognized as ideal drug delivery carriers for their biocompatibility and biodegradability and are approved by U.S Food and Drug Administration (FDA) (Blasi, 2019). A recent study reported better immunogenicity and protective efficacy of inactivated Newcastle disease vaccine with PLGA NPs in chickens (Ananda Kumar et al., 2023). Further, PLGA coated tuberculosis DNA vaccine has also shown better efficacy than the naked DNA vaccine (Mollenkopf et al., 2004). In the earlier studies, the recombinant L7/L12 protein with PLGA NPs has been shown to induce strong CMI and conferred protection against *Brucella* infection in mice (Singh et al., 2015), although the protective efficacy was shown to be lower than the live attenuated *B. abortus* S19 vaccine. The main disadvantage with PLGA NPs is the burst release at initial stage, which can be overcome by coating with chitosan that can prolong the drug delivery (Wang et al., 2013). Chitosan (CS) is a polysaccharide and copolymer of glucosamine and N-acetyl glucosamine, derived from chitin which is found in shells of crustacean (Darwesh et al., 2018). It is also biodegradable, highly biocompatible, non-toxic with immunomodulatory properties (Abourehab et al., 2022) and its use has been approved by FDA (Kantak and Bharate, 2022). The CS has been used as a successful adjuvant with influenza (Ghendon et al., 2008), hepatitis A (AbdelAllah et al., 2020), rift valley fever (El-Sissi et al., 2020), Newcastle disease (Yang et al., 2020). It induces both Th1 and Th2 mediated immune response (Shim et al., 2020). Recently, our chitosan coated PLGA nanoparticle with infectious bursal disease virus provided better protection than commercial vaccine (Kumar et al., 2025). With this background, the present study has been focused towards preparation and *in vitro* characterization of CS-modified PLGA NPs and evaluation of immunogenicity and protective efficacy of *Brucella* L7/L12 DNA vaccine formulation with CS-PLGA NPs in mice model for exploring an alternative strategy for vaccination against brucellosis.

## 2. Materials and Methods

### 2.1 Ethical statement

All experimental procedures in mice were approved by Institutional Animal Ethics Committee of ICAR-Indian Veterinary Research Institute and conducted in accordance with guidelines of the Committee for the Purpose of Control and Supervision of Experiments on Animals (CPCSEA). This work was reviewed and approved by Joint Director Research and Academic section of ICAR-Indian Veterinary Research Institute (IVRI).

### 2.2 Experimental mice and the bacterial strains

Apparently healthy 6–8-week-old Swiss albino mice were obtained from the laboratory animal resource section of Indian Veterinary Research Institute (IVRI), Izatnagar, India. The mice were maintained in cages and were provided with sterile feed and water *ad libitum*. *Brucella abortus* 544 virulent strain and *Brucella abortus* S-19 live vaccine strain were obtained from the *Brucella* laboratory, Division of Veterinary Public Health, IVRI, Izatnagar.

### 2.3 Preparation of chitosan coated PLGA NPs

PLGA NPs were prepared by the solvent displacement method (Messai et al., 2005). Briefly, 0.5% (w/v) PLGA was prepared in 10 mL of acetone and added drop-wise to 10 mL of sterile filtered distilled water containing 1% polyvinyl alcohol (PVA) as emulsion stabilizer with continuous stirring. The acetone was allowed to evaporate overnight. The NPs were pelleted by high-speed centrifugation at 24,000 rpm for 30 min and resuspended in limited amount of filtered pure distilled water and stored at 4^°^C. The PLGA NPs were negatively charged and to make them cationic, chitosan was coated on preformed PLGA NPs as per previous method (Messai et al., 2005). The final coating procedure was done by incubating the preformed NPs with different concentrations of chitosan (125 µg, 250 µg, and 500 µg) per 5 mg of NPs in 1 mL acetate buffer (pH 5.5) with continuous shaking for 1 h at room temperature (25^°^C). The CS-PLGA NPs were pelleted by centrifugation at 20000 rpm for 10 min. The coating of the chitosan was confirmed by analyzing zeta potential of the NPs and by transmission electron microscopy (TEM). For cellular uptake study, PLGA NPs encapsulating the fluorescent dye (coumarin-6) were prepared by solvent evaporation method (Vila et al., 2002).

### 2.4 Particle size and zeta potential analysis

The size distribution of PLGA NPs was determined by quasi-elastic light scattering at 25^°^C, at an angle of 90^°^, using a zeta sizer 3000 HS (Malvern Instruments, UK). Highly diluted colloidal dispersions of NPs in pure distilled water were used for size and zeta potential measurement. Further, the morphology and size of NPs were determined by TEM (FEI 12 BT 120 kv). The diluted NPs sample was loaded on copper grids stained with 2% uranyl acetate and observed under transmission electron microscope.

### 2.5 Cytotoxicity study of cationic PLGA NPs

Vero cells were seeded in 96-well tissue culture plate (100 µL/well) at the concentration of 10^6^ cells/mL in DMEM complete medium, supplemented with 10% fetal bovine serum (FBS) and incubated at 37^°^C with 5% CO_2_. After attaining 80% confluency, the cells were further incubated with 100 µL of different concentrations (10, 50, and 500 µg/mL) of PLGA NPs for 24 h at 37^°^C. The negative control with 100 µL/well of medium alone and positive control with 0.2% Triton-X 100 (for cytotoxicity) were also included. After incubation the medium was removed and cells were washed twice with PBS. Then, 85 µL of fresh culture medium and 15 µL of 3-[4,5-dimethylthiazol-2-yl]-2,5-diphenyl tetrazolium bromide (MTT) dye solution (5 mg/mL in PBS) was added to the wells and further incubated for 4 h. Then, 100 µL of dimethyl sulfoxide (DMSO) was added to each well to solubilize the formazan precipitates formed and finally, optical density (OD) was measured by the microplate reader at 550 nm. The percentage of relative growth was calculated by using the formula: (OD of nanoparticle treated sample/OD of negative control) x100.

### 2.6 Cellular uptake of PLGA NPs

Murine macrophage cell line (RAW 264.7) and Vero cells were used for cellular uptake assay of NPs as per previous method (Davda and Labhasetwar, 2002). Both cell lines were maintained in DMEM (Gibco), supplemented with 10% FBS, penicillin (100 IU/mL), streptomycin (50 µg/mL) and 25 mM HEPES. The cells were seeded at 2 ×10^4^ cells/well on the cover-slips placed in a 6-well tissue culture plate. After 80% confluency, the culture medium was replaced with fresh medium without serum, containing 100 µg/mL of coumarin-6 (dye)-encapsulated PLGA NPs and incubated at 37^°^C. At different time points (30 min, 1 h, 2 h and 4 h) the cells were harvested, cover-slips were washed thrice with sterile PBS and fixed with 1:1 methanol and acetone for 15 min. The cover-slips were subsequently mounted with 90% glycerol in PBS and observed under Confocal Microscope.

### 2.7 Plasmid DNA adsorption onto CS-PLGA NPs

Plasmid DNA (L7/L12) was adsorbed onto the surface of cationic CS-PLGA NPs as per previous method (Messai et al., 2005). Briefly, 1 mg aliquots of nanoparticles (NPs) coated with different concentrations of chitosan (section 2.2) were separately dispersed in 1 mL of acetate buffer in Eppendorf tubes. These were then mixed with varying concentrations of plasmid DNA (3–18 µg/mL) and incubated on a magnetic stirrer at 4°C for 6 hours by placing the tubes in a water-filled beaker. DNA binding to NPs was checked by agarose gel electrophoresis (1% agarose, 0.5 µg /mL ethidium bromide in Tris-acetate EDTA buffer) based on mobility of the DNA-NPs complex and the free form of DNA, visualized under UV. The amount of plasmid DNA adsorbed onto NPs was derived by determining the amount of free plasmid concentration in the supernatant recovered after centrifugation (20000 rpm, 15 min), by UV spectrophotometry.

### 2.8 *In vitro* release of plasmid DNA from PLGA NPs

10 mg of plasmid DNA loaded PLGA NPs were suspended in 1 mL of PBS and incubated in shaking incubator at 37^°^C. At different time intervals (24 h, 48 h, 7 d, 14 d, 21 d and 30 d), the PBS was collected after centrifugation at 12000 rpm for 10 min and each time NPs were resuspended with 1 mL of fresh PBS. The amount of DNA release was determined spectrophotometrically by taking absorbance at 260 nm and purity of the released DNA was checked by 1% agarose gel electrophoresis.

### 2.9 *In vitro* transfection of DNA loaded CS-PLGA NPs

The RAW 264.7 cells were grown on cover-slips in 6-well tissue culture plate (10^6^ cells/well) with complete DMEM incubated at 37^°^C with 5% CO_2_. After attaining 80% confluency the medium was removed and the cells were incubated with either 10 µg of CS-PLGA+DNA loaded NPs or commercial transfection reagent lipofectamine (Invitrogen) with 10 µg DNA or naked plasmid in 0.5 mL of serum-free medium for 4 h at 37^°^C. Then, 1.5 mL of complete fresh medium was added and further incubated for 48 h at 37^°^C. The expression of transfected DNA was determined by detecting the L7/L12 protein in the cells by indirect immunofluorescence test (IIFT). Briefly, the cells grown on cover-slip were washed with PBS and fixed with chilled 1:1 methanol and acetone for 15 min. After fixation cells were again washed with PBS and blocked with 2% BSA. Then primary antibody (anti-rL7/L12 protein, raised in rabbit) at the dilution of 1:200 was added and incubated at 37^°^C for 1 h. Subsequently, cover-slips were washed with PBS and incubated with secondary antibody, goat anti-rabbit FITC (Sigma) at the dilution of 1:100 for 1 h at 37^◦^C. Finally, the cover-slips were washed with PBS and removed from the plate and mounted with 90% glycerol in PBS on clean glass slides and observed under Confocal Laser Scanning Microscope.

### 2.10 Immunization experiment

Seventy-five healthy Swiss albino mice were randomly divided into five groups (n=15 per group) and immunized intramuscularly in the left tibialis anterior muscle with various formulations (Table 1) in 100 µL of phosphate buffered saline (PBS).

**Table 1.** Experimental design and immunization schedule. All the groups were boosted 3 weeks post priming.

| Groups<br>(n=15) | Inoculum | Dose | Route |
| --- | --- | --- | --- |
| I | Chitosan coated PLGA nanoparticles with L7/L12 DNA | 50 µg | I/M |
| II | L7/L12 naked DNA | 50 µg | I/M |
| III | Vector (control) | 50 µg | I/M |
| IV | PBS (negative control) | 100 µl | I/M |
| V | *Brucella abortus* S19 vaccine | 1 x 10<sup>5</sup> | I/P |
|  | (positive control) | CFU/mice |  |

### 2.11 Serum antibody levels

The levels of L7/L12 antigen specific total IgG/IgG1 and IgG2a antibodies in the serum samples of vaccinated and control groups at two weeks post priming, two weeks post booster and two weeks post-challenge were measured by indirect ELISA. Briefly, 96 well ELISA plates (Nunc) were coated with 100 µL of purified L7/L12 antigen (2.5 µg/mL) in carbonate-bicarbonate buffer (0.05M, pH 9.6) at 4^°^C by overnight incubation. The plates were washed thrice with phosphate-buffered saline containing 0.05% Tween-20 (PBS-T) and blocked with 1% BSA in PBS-T. After washing again thrice, 100 µL of the serially diluted sera samples were added and incubated for 90 min at 37°C. Then, after washing the plates were incubated with 100 µL/well of goat anti-mouse IgG or IgG1or IgG2a HRPO conjugates (Sigma, USA) at 1:10,000 dilution for 1 h at 37^◦^ C. After washing, 100 µL of substrate solution containing 200 µmol of o-phenylenediamine (OPD) and 0.04% H_2_O_2_ was added to each well and incubated for 15 min at room temperature. The reaction was stopped by addition of 50 µL/well of 1 M H_2_SO_4_ and the absorbance was measured at 492 nm. The mean OD of 10 unimmunized control mice sera samples at 1:40 dilutions plus 2 times standard deviation of its mean value were taken as the cut off. The titer of each sample was calculated as the reciprocal of the highest dilution that gave OD higher than the cut off value. The experiment was performed in triplicates and repeated twice.

### 2.12 Lymphocyte proliferation assay

After 1^st^, 2^nd^ and 4^th^ week of booster immunization, three mice from each group were sacrificed and their spleens were collected under aseptic conditions for splenocytes separation to detect *in vitro* cell mediated immune response by lymphocyte proliferation assay. Several perfusions were made into spleen with sterile PBS by using 1 mL tuberculin syringe to wash out splenocytes. About 3 mL of collected splenocytes was layered over 2 mL of histopaque-1077 (Sigma, USA) and centrifuged at 1600 rpm for 40 min at 25^◦^C (Chaturvedi et al., 2010). The interface ring rich with mononuclear cells were collected carefully and washed twice with PBS at 1200 rpm for 10 min. Finally, the cells were suspended in phenol red free RPMI-1640 complete medium and 100 µL of cell suspension (5 × 10^6^ cells /mL) were added to triplicate wells in 96 well flat-bottom tissue culture plates. The splenocytes were stimulated with 100 µL/well of either purified rL7/L12 protein (10 µg/mL) or Con A (20 µg/mL) or medium alone (unstimulated control) by incubating at 37^◦^C with 5% CO_2_. After 96 h, 100 µL of supernatant was carefully removed and 15 µL of MTT solution (5 mg/mL in PBS) was added to each well and further incubated for 4 h at 37^◦^C. Finally, 150 µL of DMSO was added to each well and mixed thoroughly in order to dissolve formazan crystals. The intensity of colour development was measured by taking absorbance at 550 nm in a microplate reader. Stimulation index (S.I) was calculated as the ratio between mean OD value of stimulated cells and the mean OD value of the unstimulated control.

### 2.13 Cytokine assay by Sandwich ELISA

Splenocytes of three mice from each group were isolated and plated in 96-well flat bottom tissue culture plates as described (section 2.11). Cells were stimulated with either rL7/L12 protein (10 µg/mL) or Con A (20 µg/mL) or media alone (unstimulated control) for 48 h at 37^◦^C with 5% CO_2_. After stimulation, supernatant from all the wells were collected in sterile eppendorf tubes for cytokine assay. Quantitation of murine IFN-γ, IL-2, and IL-4 was done by using Sandwich ELISA kit (Cytolab/ Pepro Tech Inc.) as per instructions in the kit.

### 2.14 Protective efficacy

Four weeks after the booster immunization, all the mice were challenged intraperitoneally with 10^5^ colony forming units (CFU) of *Brucella abortus*, strain 544 in 200 µL of sterile PBS. Four weeks after challenge, five mice from each group were sacrificed for bacterial clearance assay in spleen. The spleen was removed aseptically and homogenized in 2 mL of tryptose phosphate broth (TPB). 20 µL of suspension was placed on TP agar, incubated for 96 h at 37^◦^C and *B. abortus* 544 colonies were counted. The difference of log organism burden between the PBS control group and each vaccinated group was considered as protection in terms of bacterial clearance. At the end of experiment all mice were humanely euthanized in a CO_2_ chamber.

### 2.15 Statistical analysis

Data are presented as mean ± standard error of mean. The appropriate statistical test used are indicated in the corresponding figure legends. *P* values less than 0.05 were considered as statistically significant. (* *P* < 0.05; ** *P* < 0.01; *** *P* < 0.001 and **** *P* < 0.0001). GraphPad Prism version 8.0.2 (San Diego, CA, USA) was used for data analysis.

## 3. Results

### 3.1 Characterization of CS-PLGA NPs

On zeta potential analysis, the blank PLGA NPs showed negative charge of −5 mV. The surface of PLGA NPs was modified by adsorbing chitosan (cationic polymer) to facilitate binding of negatively charged DNA. The chitosan coated PLGA NPs were spherical in shape with almost homogeneous size distribution, with mean size of 165 nm as determined by particle size analyzer and by TEM. The coating of chitosan polymer on PLGA NPs was indicated by observation of two layers consisting of PLGA core surrounded by a thin layer of chitosan under TEM. The chitosan coating was further confirmed by the zeta potential of CS-PLGA NPs, which exhibited positive charge (+20 mV) in distilled water.

### 3.2 Cytotoxicity study of cationic PLGA NPs

Cytotoxic effect of cationic PLGA NPs (if any) on Vero cells was examined by MTT assay. Different concentrations of cationic PLGA NPs (10 µg, 50 µg and 500 µg/mL), when incubated with cells showed 98%, 97% and 95% relative growth, respectively, indicating that cationic NPs were non-toxic, whereas the positive control with Triton-X (0.2%) showed relative growth below 10%.

### 3.3 Cellular uptake of PLGA NPs

A time dependent cellular uptake of the coumarin-6 fluorescent dye-encapsulated PLGA NPs was observed in both RAW 264.7 and Vero cells. The confocal microscopic images revealed that after 30 min of incubation NPs were distributed at the periphery of the cytoplasm and as the time of incubation increased from 1 h to 4 h, there was increase in intensity of fluorescence and the number of cells showing fluorescence in all over the cytoplasm and the nucleus of the cells (Fig. 1A, B). The cellular uptake of PLGA NPs was quicker and more efficient in RAW 264.7 cells as compared to non phagocytic Vero cells at various time points. The z-stack image of cells by confocal microscopy showed internal localization of NPs at 4 h of incubation.

**Fig. 1.**
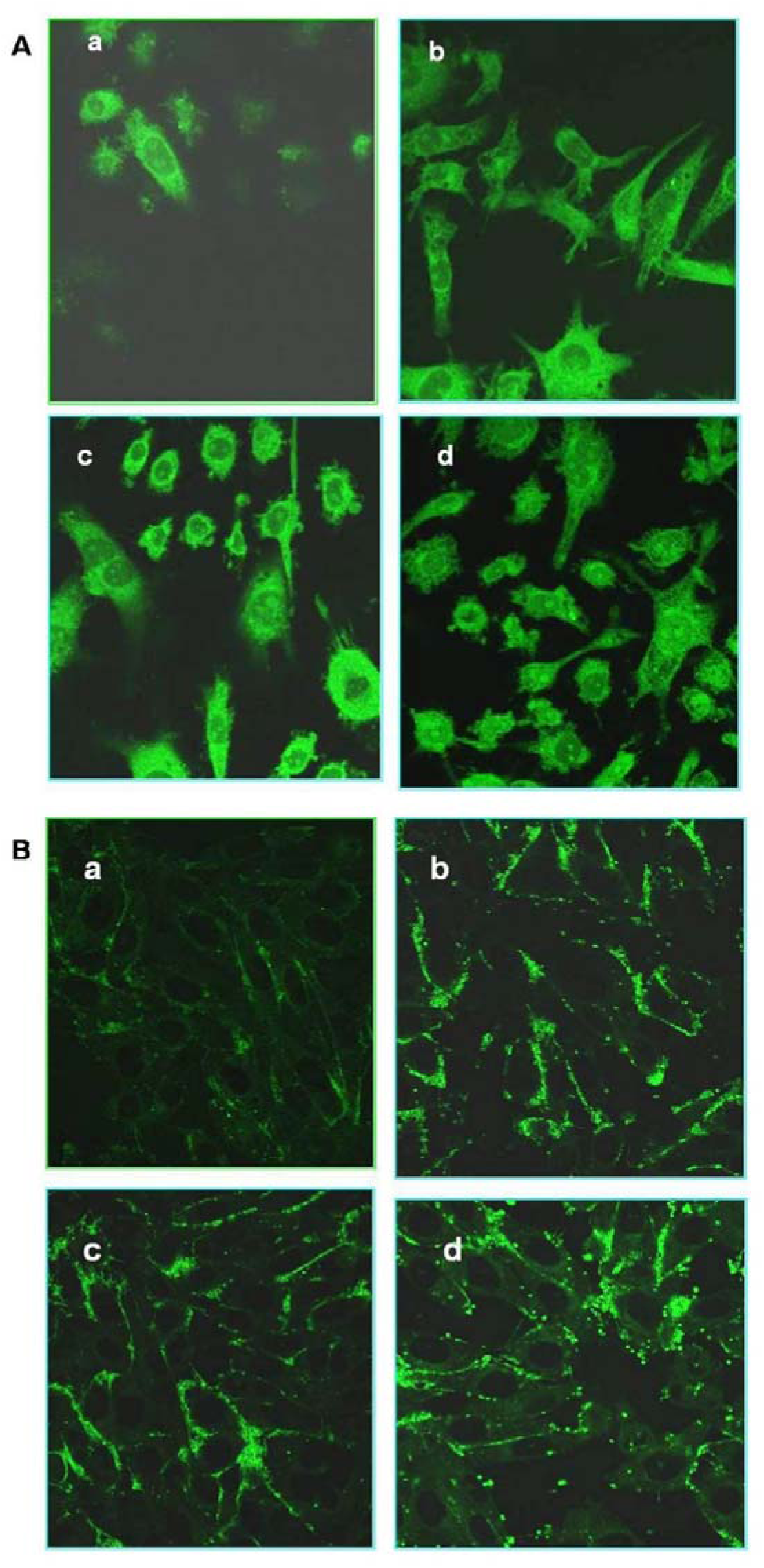
Confocal microscopic images of cellular uptake of coumarin-6 dye encapsulated PLGA NPs in murine macrophage (RAW 264.7) cell line (1A) and in non phagocytic Vero cells (1B). Cells were imaged at 30 min (a), 1 h (b), 2 h (c) and 4 h (d) post adsorption. Images were obtained by FITC filter showing time dependent increase in the intensity of fluorescence in cytoplasm and nucleus

### 3.4 Plasmid DNA adsorption to CS-PLGA NPs

Polyanionic L7/L12 DNA binding with the cationic CS-PLGA NPs was confirmed by immobilization of the DNA-NPs complex in agarose gel. The optimum DNA binding (12 µg of DNA per 1 mg of CS-PLGA NPs) was observed, when PLGA NPs were coated with 125 µg/mL of chitosan (section 2.2), with 90-100% binding efficiency and 1.2% loading efficiency. Further, DNA adsorption resulted in a shift of the charge of CS-PLGA NPs to negative zeta potential −45 mV.

### 3.5 *In vitro* DNA release kinetics

The *in vitro* release rate of plasmid DNA from the cationic PLGA NPs were initially about 25% within 24 h in PBS at 37^◦^C. Subsequently, the rate of release was slower, but up to 30 d of incubation about 90% of the DNA was released. The DNA released at different time points i.e. 24 h, 48 h, 7 d, 14 d, 21 d and 30 d were intact and showed single band on electrophoresis on 1% agarose gel indicating stability of DNA even after adsorption onto PLGA NPs.

### 3.6 *In vitro* transfection of DNA loaded CS-PLGA NPs

After *in vitro* transfection, the localization of expressed protein in RAW 264.7 cells were detected by IIFT using specific antiserum. The cells transfected with L7/L12 DNA loaded CS-PLGA NPs and commercial reagent lipofectamine with DNA showed fluorescence, indicating transfection and subsequent expression of specific protein in the cells. However, commercial transfection reagent (lipofectamine) showed a higher number of transfected cells as compared to the plasmid DNA loaded CS-PLGA NPs (Fig.2).

**Fig. 2.**
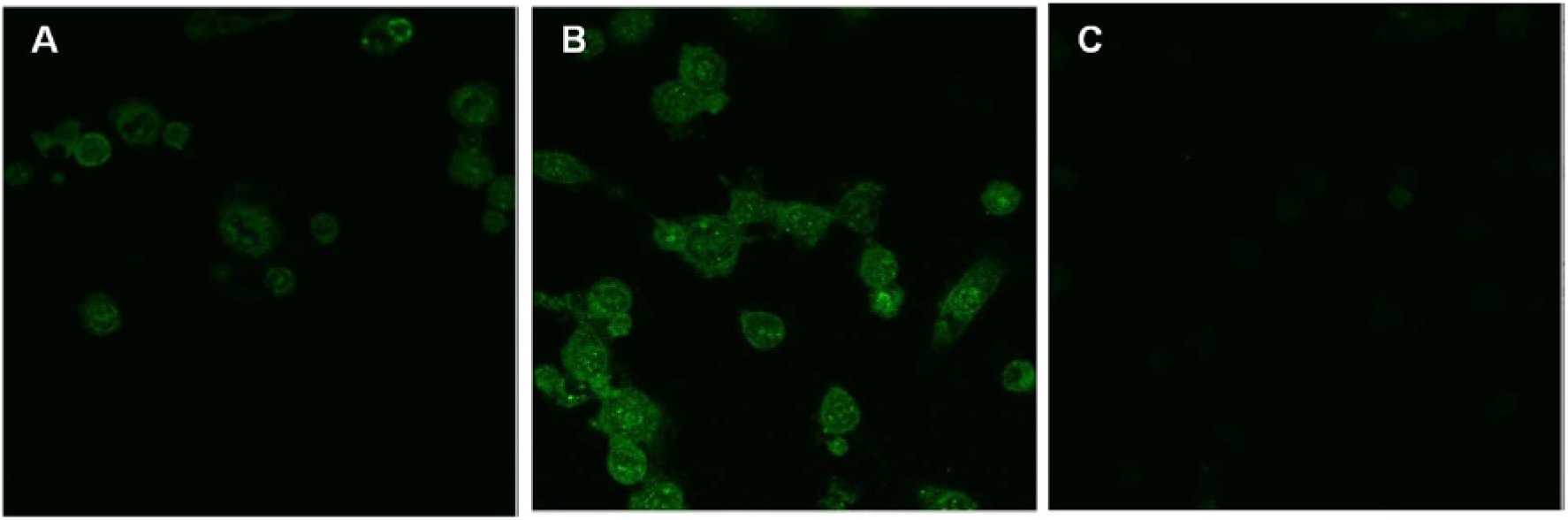
Confocal microscopic images of murine macrophage (RAW 264.7 cells) transfected with (A) L7/L12 plasmid DNA loaded CS-PLGA NPs, (B) plasmid DNA complexed with lipofectamine showing *in vitro* protein expression, detected by IIFT and (C) cells transfected with naked plasmid DNA as control

### 3.7 Serum antibody levels in immunized mice

The levels of IgG, IgG1 and IgG2a antibodies at 14 d post priming and 14 d post booster immunization as well as at two weeks post challenge infection with *B. abortus* are shown in Fig. 3A, 3B, 3C, respectively. There was anamnestic response observed after booster and post challenge with the maximum antibody titer recorded after challenge infection. The antibody titers of IgG in group-I (DNA+NPs) and group-II (naked DNA) after booster immunization were 750 and 450, respectively. Similarly, the titers of IgG1 were 113 and 75, and IgG2a were 225 and 163, respectively. The mice, sham-immunized with vector DNA (group III) and PBS control (group IV) did not show any specific antibody response.

**Fig. 3.**
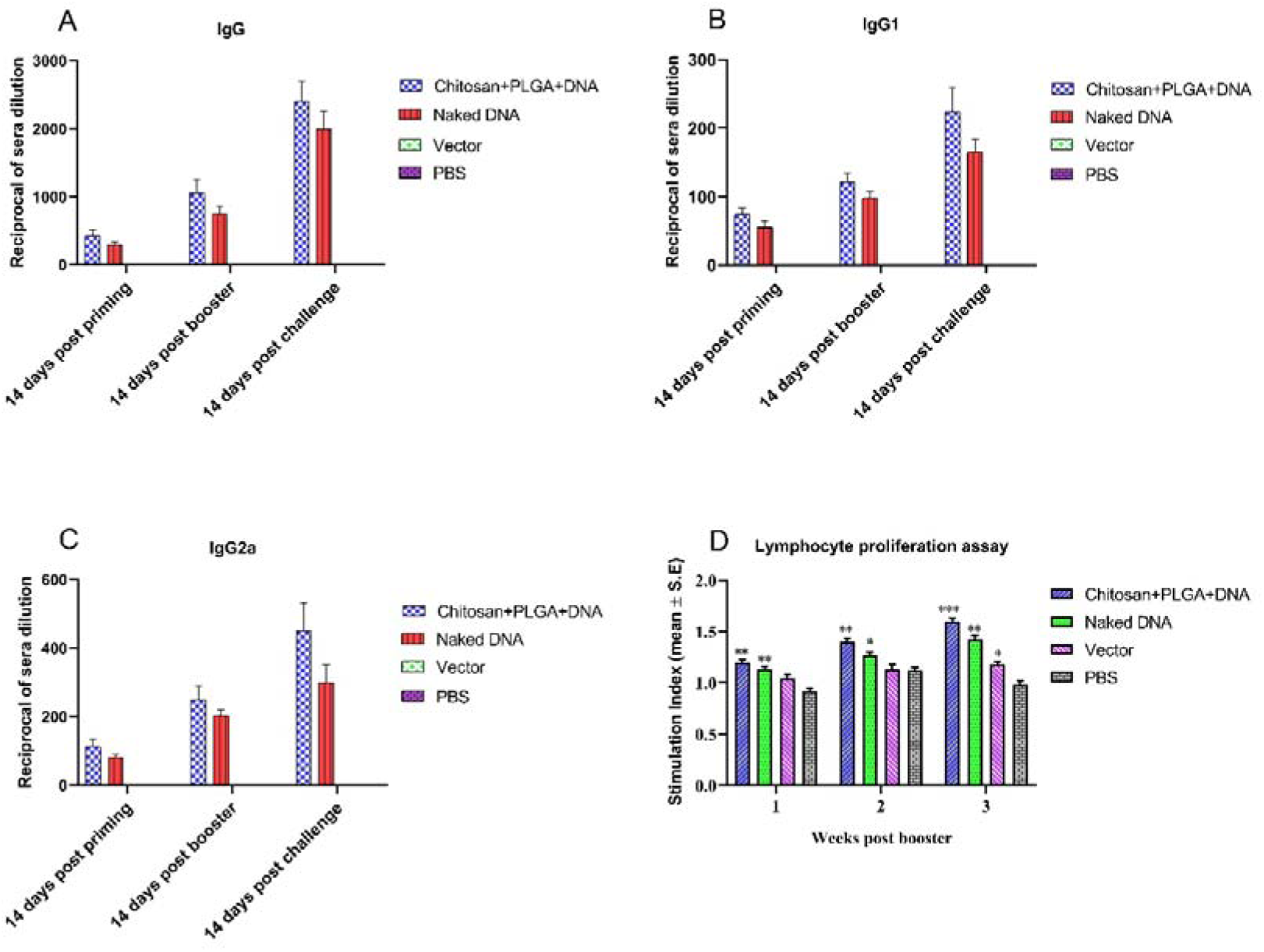
*Brucella abortus* rL7/L12 specific antibody titers; IgG (3A), IgG1 (3B) and IgG2a (3C) in the serum samples of mice injected with CS-PLGA+DNA nanoparticles or naked DNA or vector or PBS, at different time intervals, determined by indirect ELISA. Proliferation of splenocytes of mice immunized with CS-PLGA+DNA nanoparticles or naked DNA or vector or PBS, determined by MTT colorimetric assay (3D). At each time point 3 mice were sacrificed to collect splenocytes and the assay was set up in triplicates. Within weeks, each group was compared to PBS group by student’s-t test.

### 3.8 Lymphocyte proliferation

There was gradual increase in proliferation of splenocytes of the mice immunized with L7/L12 DNA loaded NPs (group I) and naked DNA (Group II) at different time points post booster immunization. Splenocytes from group I and II showed significantly higher (*P* <0.01) proliferation than that of control group. Mice of group I exhibited highest lymphocyte proliferation (1.60±0.028) followed by group II (1.42±0.043) at 4^th^ week of booster (Fig. 3D).

### 3.9 Cytokine levels

The IFN-γ levels in the culture supernatant was maximum at 4^th^ week post-booster immunization in group I (2500±0.053 pg/mL) and group II (1768±0.034 pg/mL) as compared to the control group IV (67±0.027 pg/mL) (Fig. 4A). The production of IL-2 and IL-4 upon stimulation with rL7/L12 was found to be weak as compared to IFN-γ. The group I and II showed highest amount of IL-4 (255 ±0.063 and 185±0.14 pg/mL, respectively) at 2^nd^ week post-booster. However, highest production of IL-2 was detected at the 4^th^ week in group I (419±0.078 pg/mL) and group II (349±0.045pg/mL) (Fig. 4B & 4C).

**Fig. 4.**
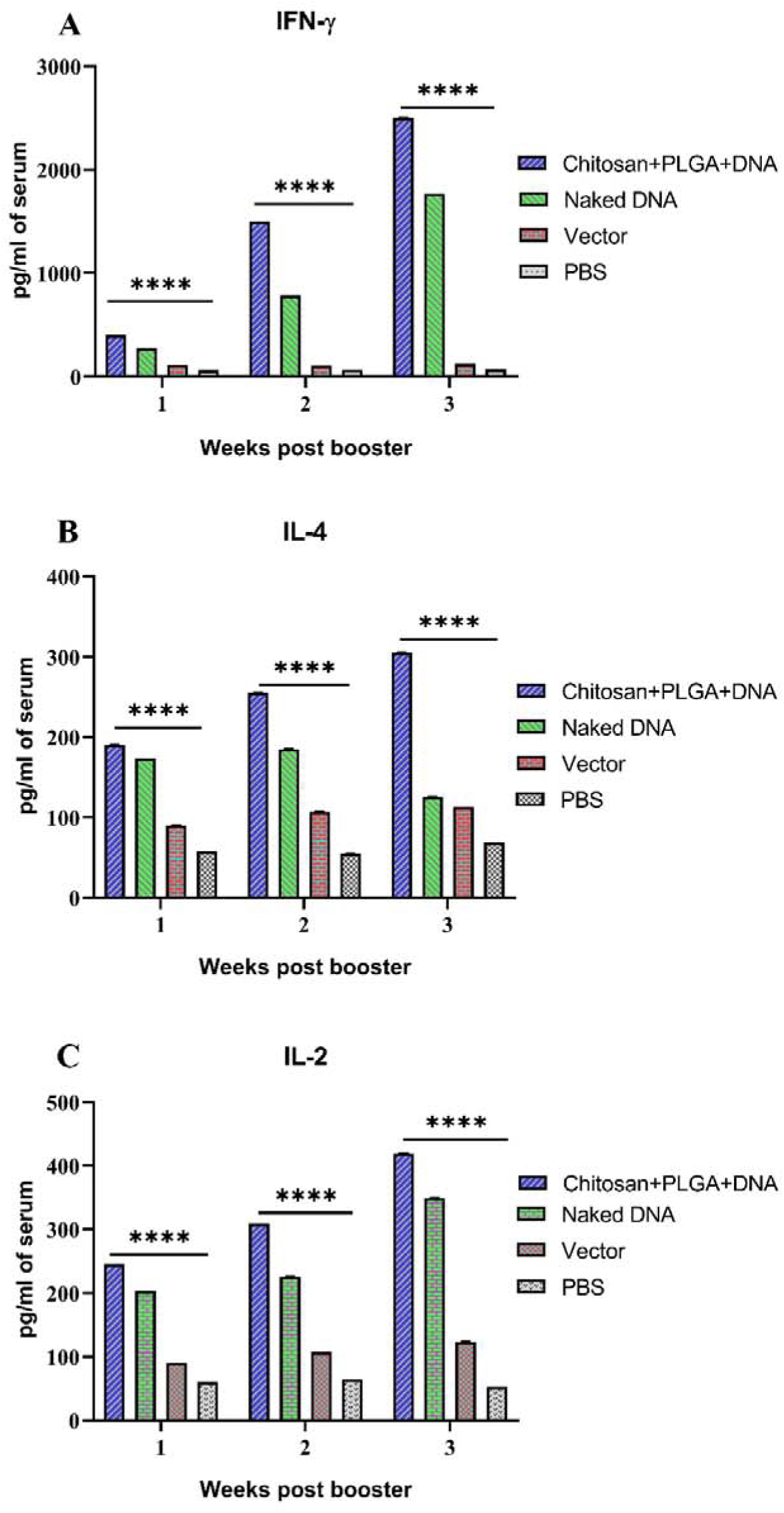
IFN-γ (4A), IL-4 (4B) and IL-2 (4C) induced by the splenocytes of different groups of immunized mice. At each time points 3 mice were sacrificed to collect splenocytes. After stimulation of splenocytes, the culture supernatants were collected and quantitation of cytokines was done by Sandwich ELISA kit. Samples were tested in triplicates. Each bar represents mean concentration of cytokine in pg/mL and the differences were analyzed within weeks by one-way ANOVA.

### 3.10 Protection against challenge infection

Following challenge infection in the vaccinated mice with virulent *B. abortus* 544, the difference in log organism burden between the PBS control group (unimmunized mice) and each of the vaccinated groups was considered as protection in terms of reduction in bacterial load (Fig. 5). The log protection in group I (DNA+NPs), group II (naked DNA) and group V (*B. abortus* S19 vaccine) was 1.9, 1.2 and 2.37, respectively (Table 2). These findings demonstrate group I (DNA+NPs, *P*<0.0001), group II (naked DNA, *P*=0.0017) and group V (S-19 vaccine, *P* <0.0001) showed significantly higher protection as compared to the PBS control group.

**Fig. 5.**
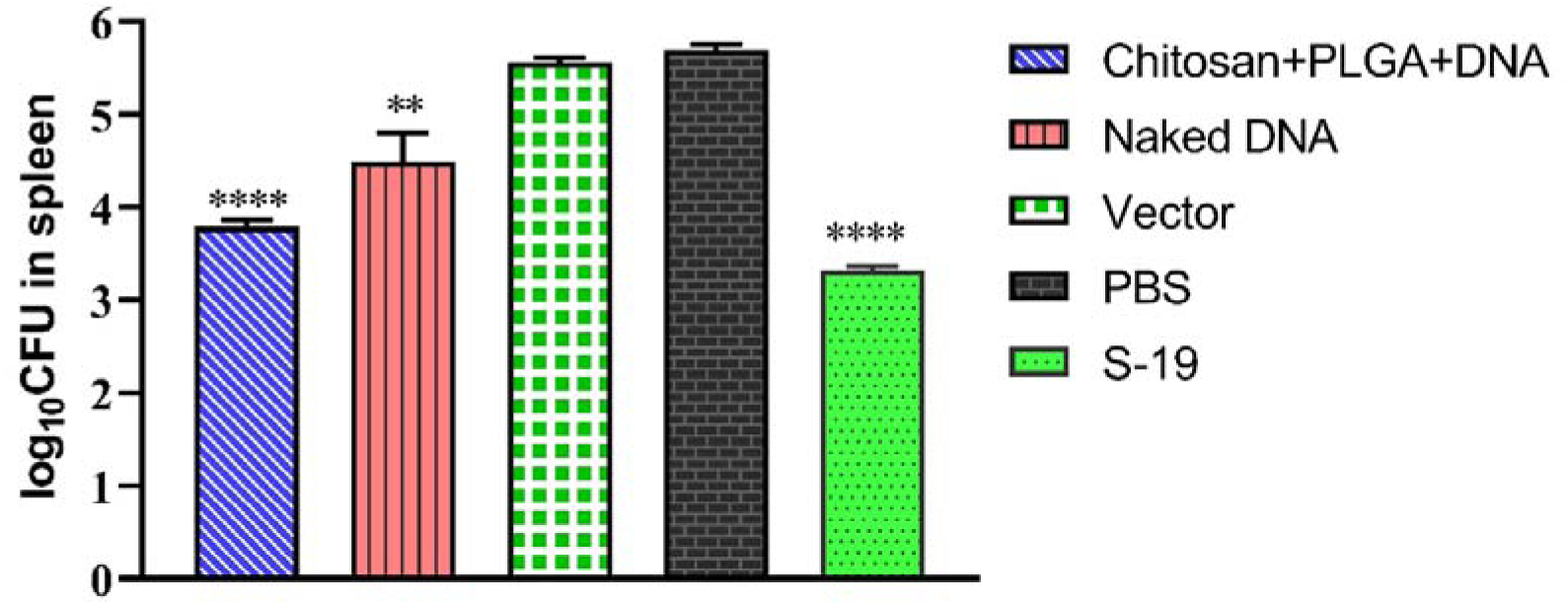
The mean *Brucella* copies (CFU) in the spleen of immunized and control groups of mice were calculated on 28 d post-challenge with virulent *Brucella abortus* 544. The data represent the mean log_10_ CFU± standard error. Each group was compared to PBS group by student’s-t test.

**Table 2.** Bacterial loads in the spleen of different immunized groups of mice after challenge infection. Four weeks after the booster immunization, all the mice were challenged intraperitoneally with 10^5^ CFU of *Brucella abortus*, strain 544 and then organism burden in the spleen was determined four weeks post challenge. Units of protection was calculated by subtracting mean log_10_ CFU in the spleen of vaccinated groups from that of PBS control group and differences of *Brucella* organism counts were analyzed by student’s-t test. Values marked with a (*P* <0.0001) and b (*P* <0.001) significantly differ from PBS control

| Experimental groups | Log<sub>10</sub> CFU of *B. abortus* in spleen (mean±SE) | Log<sub>10</sub> units of protection |
| --- | --- | --- |
| **I.** Chitosan coated PLGA nanoparticles with L7/L12 DNA | 3.79±0.07<sup>a</sup> | 1.9 |
| **II.** Naked DNA | 4.49±0.31<sup>b</sup> | 1.2 |
| **III.** Vector | 5.56±0.054 | 0.13 |
| **IV.** PBS | 5.69±0.067 | 0.00 |
| **V.** *B. abortus*-S19 vaccine | 3.32±0.049<sup>a</sup> | 2.37 |

## 4. Discussion

Brucellosis is an important zoonotic disease that mostly affects cattle and many cases in humans have also been reported. As there is no availability of licensed human vaccine, controlling the disease in animals might reduce the spread of infection to humans. Although few vaccines are available to control the disease in animals using *B. abortus* S-19, RB-51 and Rev-1 strains etc., these live vaccines are pathogenic to humans and induce abortion in pregnant cattle. These drawbacks highlight the need for improved vaccine strategies. Hence, some previous studies were focused to deliver safe vaccine that can confront the disadvantages associated with the existing vaccines, but there was limited success and none of them was better than the standard live vaccines S-19 or RB-51. However, an alternate strategic approach using DNA vaccines was able to induce both humoral and cellular immune responses. Since *Brucella abortus* is an intracellular pathogen, the cell-mediated immunity plays a crucial role for control of the disease, in which, L7/L12 DNA has been reported to induce strong T cell response (Mallick et al., 2007). The recombinant L7/L12 protein and plasmid encoding the L7/L12 gene can elicit strong CMI and protection from *Brucella* infection in mice; however, the protective efficacy is much lower than the live attenuated *Brucella abortus* vaccine S19 (Kurar, 1997; Luo et al., 2006; Yu et al., 2007; Isore et al., 2014). In earlier studies, *Brucella* genes encoding ribosomal protein L7/L12 (Senevirathne et al., 2020), divalent L7/L12-Omp 16 (Luo et al., 2006), lumazine synthase DNA (Velikovsky et al., 2002), bacterioferritin DNA (Al-Mariri et al., 2001b) have also been tested as DNA vaccines in mice.

Given the potential of DNA vaccines in inducing both humoral and CMI response, we have investigated the immunogenicity and protective efficacy of chitosan-modified PLGA NPs with L7/L12 plasmid in mice against *Brucella* infection. Our results revealed that delivery of plasmid DNA through the CS-PLGA cationic NPs was successful and further *in vitro* transfection study established the protein expression in RAW 264.7 cell line (Murine macrophage). Upon immunization the plasmid was likely expressed in host cells and stimulated immune response. The cationic NPs were optimum in size to be captured by antigen presenting cells to elicit immune response as evidenced by our results. The CS-PLGA NPs release DNA in a controlled manner as demonstrated in our *in vitro* release kinetics, which favors prolonged immune response. In correlation to our findings, previous studies reported that PLGA-chitosan NPs provide sustained release of the drug up to 140 h *in vitro* and could be a good approach to improve bioavailability of the entrapped drug (Bandil et al., 2023). Recently, chitosan-modified PLGA NPs have also been used for drug delivery to enhance gastric retention and controlled release (Lestari et al., 2024). Our immunization experiment revealed that the DNA loaded CS-PLGA NPs have induced significant levels of antibody as well as CMI response. Further, this experimental vaccine formulation afforded protection in terms of bacterial clearance in the spleen up to 1.9 log_10_. Previous studies reported that vaccine candidates (L7/L12 protein) generated a log protection of 1.69 (Singh et al., 2015), 1.61 (Mallick et al., 2007) and the L7/L12 DNA vaccine showed 1.26 (Kurar, 1997). In correlation with previous reports, the higher protection level observed in the present study (1.9 logLL units) may be attributed to the use of CS-PLGA nanoparticles, which favors in sustained release and the adjuvant effect helps in potent induction of humoral as well as CMI response that ultimately conferred efficient protection against virulent *Brucella* organisms. Earlier workers also have reported that chitosan was able to enhance neutralizing antibody titers against rift valley fever virus and pertussis toxin (Jabbal-Gill et al., 1998; El-Sissi et al., 2020). Further, the induction of cytokines is the prime criterion for generating a potent immune response. In our study, cytokine profiling revealed that the L7/L12 loaded NPs elicited a significant expression of IFN-γ in the immunized mice (Fig. 4A), which might have played a crucial role in the activation of macrophages and cytotoxic T-lymphocytes (CTL) and facilitated the clearance of bacteria. The earlier workers also reported that the IFN-γ plays an important role in the resolution of brucellosis (Zhan and Cheers, 1993; Eze et al., 2000). In addition, the CS-PLGA NPs with L7/L12 DNA vaccine has stimulated strong T cell response than the L7/L12 DNA alone as evidenced by antigen specific lymphocyte proliferation (Fig.3D). The earlier worker also reported that intramuscular injection of the L7/L12 gene resulted in specific antibodies and T-cell responses with significant protection against *B. abortus* infection (Kurar, 1997). Similarly, a bivalent DNA vaccine encoding the Omp16 and L7/L12 genes induced strong humoral and cellular immune responses with significant protection against *B. abortus* 544 (Luo et al., 2006).

The IgG2a is essential for protection against intracellular pathogens (Rostamian et al., 2017). The enhanced titer of IgG2a over IgG1 in the present study indicated the isotype switching under the influence of IFN-γ (Th1 cytokine) due to a predominant stimulation of cellular response. In correlation with the present findings, earlier workers also reported an induction of IgG2a by the *Brucella* DNA vaccine encoding lumazine synthase and bacterioferritin/P39 (Al-Mariri et al., 2001a; Velikovsky et al., 2002). Cationic microparticles adsorbed with plasmid DNA encoding Japanese encephalitis virus envelope protein also induced IgG2a over IgG1 (Kaur et al., 2004). Induction of strong IFN-γ along with IgG2a indicates stimulation of Th1 biased cell mediated immunity. In the present study, production of IL-4 by lymphocytes of the immunized mice indicating its role in humoral immunity and IL-2 production supports the CMI response. In our study following challenge, an anamnestic response in the antibody titer and cytokines levels were observed. The similar finding of post-challenge enhancement in antibody titer has been reported in avian influenza vaccine experimentation (Panickan et al., 2022). Overall, the L7/L12 DNA loaded CS-PLGA NPs elicited efficient humoral and cell-mediated immune response along with better protection against *Brucella* infection. Further, unlike live vaccines the L7/L12 DNA will be safe as it is a subunit vaccine that lacks virulence factor. Therefore, vaccines containing recombinant protein or their coding DNA can be suitable alternatives to the conventional vaccines, because they can reduce unwanted side effects. To the best of our knowledge, this is the first study on chitosan-modified PLGA NPs with L7/L12 DNA vaccine in mice against *Brucella* infection.

## 5. Conclusion

The plasmid DNA encoding *B. abortus* L7/L12 with the CS-PLGA NPs elicited both cell mediated and humoral immune response in the immunized mice and also afforded protection against the virulent *Brucella abortus* 544 challenge infection. The study indicated that the NPs made up of a combination of PLGA and chitosan is a promising DNA delivery system that is more effective than the traditional adjuvant and also provides an insight for the development of effective vaccines against *Brucella*. The present findings warrant further investigation of the mechanistic activity and the potential benefits of PLGA-chitosan NPs based delivery system cum adjuvant.

## Acknowledgement

The authors are thankful to Director, ICAR-IVRI for arranging necessary facilities and funding.

## Ethics approval

Approved by Institutional Animal Ethical Committee

## Availability of data and materials

All data are included in the manuscript

## Authors’ contributions

S.D and U.K.C conceptualized and designed the study, performed experiments and wrote the manuscript. S.P. performed data analysis and prepared figures. V.K and P.M.S edited and reviewed the manuscript. All the authors read and approved the final manuscript.

